# Insulin inhibits protein phosphatase 2A to impair β-adrenergic receptor function

**DOI:** 10.1101/2020.09.17.302448

**Authors:** Anita Sahu, Yu Sun, Sromona Mukherjee, Conner Witherow, Kate Stenson, John J.G. Tesmer, Maradumane L. Mohan, Sathyamangla V. Naga Prasad

**Affiliations:** Department of Cardiovascular and Metabolic Sciences, Lerner Research Institute, Cleveland Clinic, Cleveland, OH, 44195

## Abstract

Insulin impairs β2-adrenergic receptor (β2AR) function through G protein-coupled receptor kinase 2 (GRK2) by phosphorylation but less is known about dephosphorylation mechanisms mediated by protein phosphatase 2A (PP2A). Pharmacologic or genetic inhibition of phosphoinositide 3-kinase γ (PI3Kγ) unexpectedly resulted in significant reduction of insulin-mediated β2AR phosphorylation. Interestingly, β2AR-associated phosphatase activity was inhibited by insulin but was reversed by knock-down of PI3Kγ showing negative regulation of PP2A by PI3Kγ. Co-immunoprecipitation and surface plasmon resonance studies using purified proteins showed that GRK2 and PI3Kγ form a complex and could be recruited to β2ARs as GRK2 interacts with insulin receptor substrate following insulin treatment. Consistently, β-blocker pretreatment did not reduce insulin-mediated β2AR phosphorylation indicating agonist- and Gβγ-independent non-canonical regulation of receptor function. Mechanistically, PI3Kγ inhibits PP2A activity at the βAR complex by phosphorylating an intracellular inhibitor of PP2A (I2PP2A). Knock-down or CRISPR ablation of endogenous I2PP2A unlocked PP2A inhibition mediating β2AR dephosphorylation showing an unappreciated acute regulation of PP2A in mediating insulin-β2AR cross-talk.

**Summary:** Insulin impairs β2-adrenergic receptor (β2AR) function through G protein-coupled receptor kinase 2 (GRK2). We show that insulin simultaneously inhibits protein phosphatase 2A (PP2A) sustaining β2AR functional impairment. Unexpectedly, releasing PP2A inhibition by PI3Kγ preserves β2AR function despite intact insulin-driven GRK2-mechanisms.

## Introduction

Insulin binds to insulin receptor (IR) leading to phosphorylation of insulin receptor substrate (IRS) which mediates downstream signals regulating metabolism, survival and cell growth ^1^. Insulin plays a key role in energy homeostasis by regulating glucose uptake via GLUT4 membrane translocation^2^ through activation of a phosphatidylinositol 3-kinase (PI3K)-Akt pathway^3^. In contrast to this traditional role in physiology, abnormal insulin signaling underlies disease states like diabetes, obesity and the consequent metabolic syndrome. In addition, hyperinsulinemia plays an essential role in deleterious cardiac remodeling associated with hypertrophy and fibrosis leading to diabetic cardiomyopathy^4,5^. Increasing evidence also shows that conditions of hyperinsulinemia is associated with elevated sympathetic overdrive that could mediate signals through adrenergic receptors. Stimulation of β2-adrenergic receptor (β2AR) results in additional auxiliary effect on glucose uptake by phosphorylation of Akt through protein kinase A (PKA) dependent mechanisms^6^ suggesting the presence of active cross-talk between insulin and β2AR pathway.

βARs are key regulators of cardiac function as agonist-activation of the βAR results in G-protein coupling and cAMP generation leading to activation of PKA^7^. β2ARs undergo phosphorylation (desensitization) mediated by G protein-coupled receptor kinase (GRK2) that leads to β-arrestin recruitment and receptor endocytosis^7^. βAR resensitization occurs by dephosphorylation mediated by protein phosphatase 2A (PP2A) in the endosomes before recycling to the plasma membrane^8^. We have previously shown that phosphoinositide 3-kinase γ (PI3Kγ) impairs resensitization by phosphorylating the inhibitor of protein phosphatase 2A (I2PP2A) which binds to PP2A and inhibits its activity^9^. This shows that agonist-stimulation of β2ARs results in inhibition of PP2A reinforcing the desensitization of the receptor as knock down of I2PP2A reverses β2AR phosphorylation and desensitization^9^. These findings show acute regulation of β2AR function by PP2A in response to its agonist however, whether such acute regulation of PP2A exists in the cross-talk between IR and β2AR is not known.

Studies have shown functional membrane complex of IR and β2AR^10^ reflecting an active cross-talk between these two signaling systems. In contrast, to the additive effect on glucose uptake by β2AR stimulation^11,12^, insulin stimulation is shown to result in loss of β2AR signaling. The loss in β2AR signaling is measured by the inability of β2ARs to generate cAMP in response to its agonist following pre-treatment with insulin. Furthermore, studies have shown that insulin stimulation results in β2AR phosphorylation by GRK2 which is recruited to the IR-β2AR functional complex by IRS2 initiating β2AR internalization^13^. Although GRK2 non-canonically mediates desensitization of β2ARs through IRS2 recruitment, it is not known whether acute regulation/inhibition of PP2A occurs to augment insulin-mediated non-canonical β2AR phosphorylation.

Despite differential regulatory mechanisms and signaling, insulin or adrenergic stimulation results in GRK2 recruitment and PI3K activation^14^. Insulin stimulation results in recruitment of Class 1A PI3Kα through IRS whereas, β2AR activation recruits Class 1B PI3Kγ through Gβγ subunits, initiating downstream Akt activation^15^. In addition, we have previously shown that the protein kinase activity of PI3Kγ plays a key role in β2AR-mediated EGFR transactivation by phosphorylating the tyrosine kinase Src^16^. Given the role of PI3Kγ in receptor cross-talk and its function in inhibiting PP2A activity, we sought to test whether non-canonical recruitment of PI3Kγ to the IR-β2AR functional complex could inhibit PP2A activity resulting in sustained β2AR phosphorylation. Using HEK 293 cells, mouse embryonic fibroblasts and primary cardiomyocytes, we show that insulin stimulation results in significant β2AR phosphorylation which can be reversed by pharmacologic inhibition of PI3K or genetic ablation of PI3Kγ. Furthermore, we also show that PI3Kγ interacts with GRK2 to form a complex that is recruited to the IR-β2AR following insulin. Consistent with recruitment of PI3Kγ, there was significant reduction in β2AR-associated phosphatase activity following insulin which was reversed by knockdown of PI3Kγ. Moreover, CRISPR deletion of I2PP2A resulted in significant reduction in β2AR phosphorylation in spite of insulin unraveling an underappreciated role of PP2A regulation that sustains kinase-mediated function including receptor phosphorylation (desensitization).

## Results

### PI3Kγ regulates β2AR phosphorylation following Insulin stimulation

To test whether Insulin stimulation alters β2AR phosphorylation, HEK 293 cells stably expressing FLAG-β2AR (FLAG-β2AR HEK) were treated with saline (Sal), insulin (INS) or isoproterenol (ISO) as positive control. Consistent with previous studies^13^, significant β2AR phosphorylation was observed following 10 minutes of INS treatment as assessed by immunoblotting with anti-phospho-β2AR antibody [**Fig. 1A**, cumulative bar graph (n=5)]. Similarly, confocal microscopy also showed marked β2AR phosphorylation after INS (10 minutes) treatment as visualized by anti-phospho-β2AR staining (green) [**Fig. 1B, panels g & i**]. In contrast, minimal phosphorylation was observed with saline (Sal) [**Fig. 1B, panels a & c**] while ISO stimulation (positive control) showed accumulation of phosphorylated β2ARs [**Fig. 1B, panels d & f**]. Nucleus was stained by DAPI (blue). Given that ISO stimulation leads to endocytosis of β2ARs, internalization was blocked by pre-treatment of cells with sucrose and β-cyclodextrin which allows for marked membrane decoration of phosphorylated β2ARs [**Fig. 1b, panel d**]. Since it is not known whether INS stimulation results in acute internalization, these cells were not pre-treated with sucrose and β-cyclodextrin. Although INS treatment results in significant acute phosphorylation, it does not cause β2AR internalization [**Fig. 1B, panel g**]. To further determine whether INS causes β2AR phosphorylation at the membrane, FLAG-β2AR HEK cells were treated with INS and immunoblotting was performed on the plasma membrane fractions. Significant β2AR phosphorylation was observed at the plasma membranes following INS treatment compared to Sal [**Fig. 1C**, cumulative bar grant (n=4)]. To test whether endogenous β2AR also responds non-canonically to INS, β2AR phosphorylation was assessed in H9C2 cardiac myoblasts following INS stimulation. INS treatment resulted in significant β2AR phosphorylation [**Supplementary Fig. 1A**] showing conservation of non-canonical regulation of β2ARs by INS.

**Figure 1:**
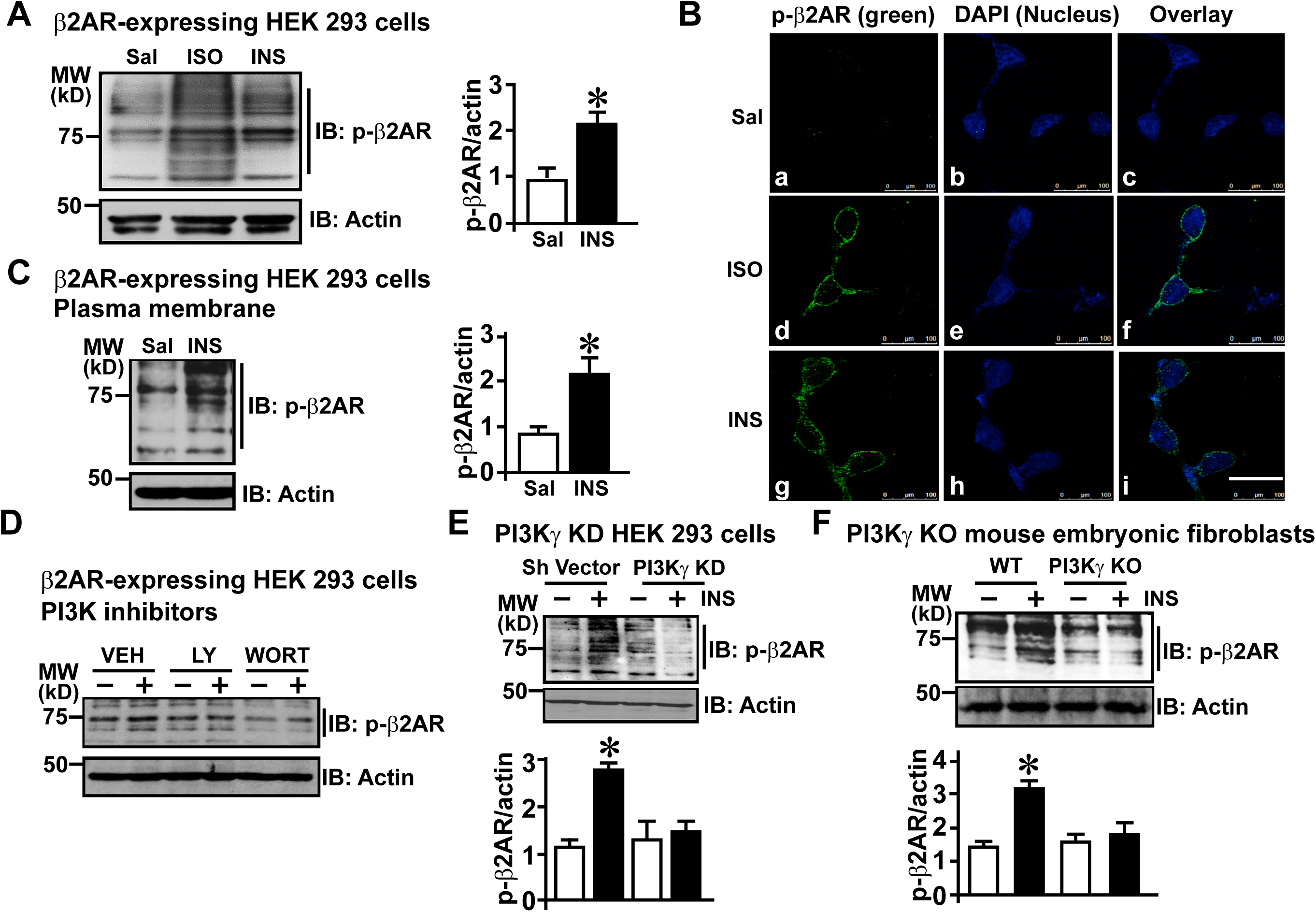
PI3Kγ regulates β2AR phosphorylation following Insulin Resistance: (A) Immunoblotting performed on cell lysates with anti-phospho-β2AR antibody to assess phosphorylation of β2AR following saline (Sal), isoproterenol (ISO, 10uM) or insulin (INS, 100nM) stimulation for 10 minutes in FLAG-β2AR HEK 293 cells. Cumulative densitometry is shown in the bar graph. (n=5) *p< 0.0001 versus control (Sal). Actin was used as loading control for all experiments. (B) FLAG-β2AR phosphorylation was visualized by confocal microscopy using anti-phospho-β2AR antibody (green) following ISO or INS stimulation for 10 minutes. Only ISO treated cells were pre-treated with internalization blockers (sucrose and β-cyclodextrin) to assess membrane decoration of stained receptors as positive control while, INS treated cells were not pre-treated with endocytosis blockers. Nucleus was visualized by DAPI (blue) staining. Scale bar: 10 μm. (C) Immunoblotting was performed on plasma membrane fraction to assess β2AR phosphorylation in FLAG-β2AR HEK 293 cells following INS stimulation for 10 minutes. Cumulative data is shown in the bar graph, (n=4) *p< 0.001 versus control. (D) Role of PI3K in INS-mediated β2AR phosphorylation was determined by pre-treatment with PI3K-specific inhibitors LY294002 (1uM) or Wortmannin (100nM). β2AR phosphorylation was assessed by immunoblotting with anti-phospho-β2AR antibody following INS stimulation for 10 minutes in HEK-FLAG-β2AR cells. (E) To specifically test the role of PI3Kγ in INS-mediated β2AR phosphorylation, HEK 293 cells with stable knock-down of PI3Kγ (PI3Kγ KD) were assessed for β2AR phosphorylation by immunoblotting with anti-phospho-β2AR antibody following the INS stimulation for 10 minutes. Cumulative data is shown in the bar graph below, (n=6), *p< 0.001 versus control (Sal). (F) Immunoblots to assess phospho-β2AR following INS stimulation for 10 minutes in mouse embryonic fibroblasts isolated from wild-type mice (WT MEF) and PI3Kγ KO mice (KO MEF). Cumulative densitometry data is shown in the bar graph, (n=5), *p< 0.001 versus control (Sal).

Since PI3Kγ inhibits β2AR-associated PP2A activity in response to traditional agonist, we tested whether PI3Kγ plays a role in regulating INS-mediated β2AR phosphorylation. FLAG-β2AR HEK cells were pre-treated with either LY294002 or Wortmannin (selective PI3K inhibitors) followed by INS treatment. Increased β2AR phosphorylation observed following INS treatment was abolished in the presence of LY294002 or Wortmannin as assessed by immunoblotting with anti-phospho-β2AR antibody [**Fig. 1D**]. Because LY294002 and Wortmannin inhibit all isoforms of PI3K, we used HEK 293 cells with stable knock-down of PI3Kγ^17^ (PI3Kγ KD) to directly test the role of PI3Kγ in INS-mediated β2AR phosphorylation. Immunoblotting using anti-PI3Kγ antibody showed significant knockdown of PI3Kγ, validating the reduction in PI3Kγ expression in the PI3Kγ KD cells compared to control vector (Sh Vector) cells [**Supplementary Fig. 1B**]. Sh vector control cells and PI3Kγ KD cells were treated with INS and β2AR phosphorylation was assessed by immunoblotting with anti-phospho-β2AR antibody. While significant β2AR phosphorylation was observed in control Sh vector cells, this was completely abrogated in PI3Kγ KD cells [**Fig. 1E, cumulative bar graph (n=6)**]. This shows that PI3Kγ plays a key role in regulating β2AR phosphorylation in response to non-canonical regulation by INS. To further strengthen our finding on PI3Kγ regulating β2AR phosphorylation by INS, we used primary mouse embryonic fibroblasts (MEFs) from PI3Kγ knockout (PI3Kγ KO) mice and wild-type (WT) control mice^18^. WT and PI3Kγ KO MEFs were stimulated with INS and endogenous β2AR phosphorylation assessed by immunoblotting. Robust β2AR phosphorylation was observed in WT MEFs which was significantly reduced in PI3Kγ KO MEFs [**Fig. 1F, cumulative bar graph (n=5)**] confirming the key role of PI3Kγ in INS-mediated β2AR phosphorylation. These results show that PI3Kγ plays a pivotal role in maintaining β2AR phosphorylation following insulin stimulation.

### Non-canonical regulation of β2AR by INS is β-arrestin- and β-blocker-independent

Given that INS causes β2AR phosphorylation, we tested whether this phosphorylation impacts β2AR response to its classical agonist ISO by measuring cAMP generation through a luciferase-based assay. A subset of FLAG-β2AR HEK cells transiently expressing cAMP luciferase were pre-treated with INS and challenged with βAR agonist ISO. ISO stimulation resulted in appreciable generation of cAMP which was significantly reduced following pre-treatment with INS [**Fig. 2A**] suggesting that INS impairs β2AR mediated cAMP generation and downstream signaling. Classically, β2AR phosphorylation results in recruitment of scaffolding protein β-arrestin that mediates receptor internalization^19^, and so we assessed whether non-canonical phosphorylation of β2AR by INS recruits β-arrestin. HEK 293 cells stably expressing β2AR and β-arrestin-2-GFP were stimulated with ISO or INS and β-arrestin recruitment to the plasma membrane was assessed by confocal microscopy. ISO stimulation resulted in recruitment of β-arrestin-GFP to the plasma membrane [**Fig. 2B, panels d & f**] as noted by significant clearance from the cytoplasm. Interestingly, no β-arrestin recruitment to the membrane was observed following INS treatment [**Fig. 2B, panels g & i**]. Saline (Sal) vehicle treatment also did not alter β-aresstin localization [**Fig. 2B, panels a & c**]. These observations show that at least in acute settings, INS mediated β2AR desensitization is β-arrestin independent.

**Figure 2:**
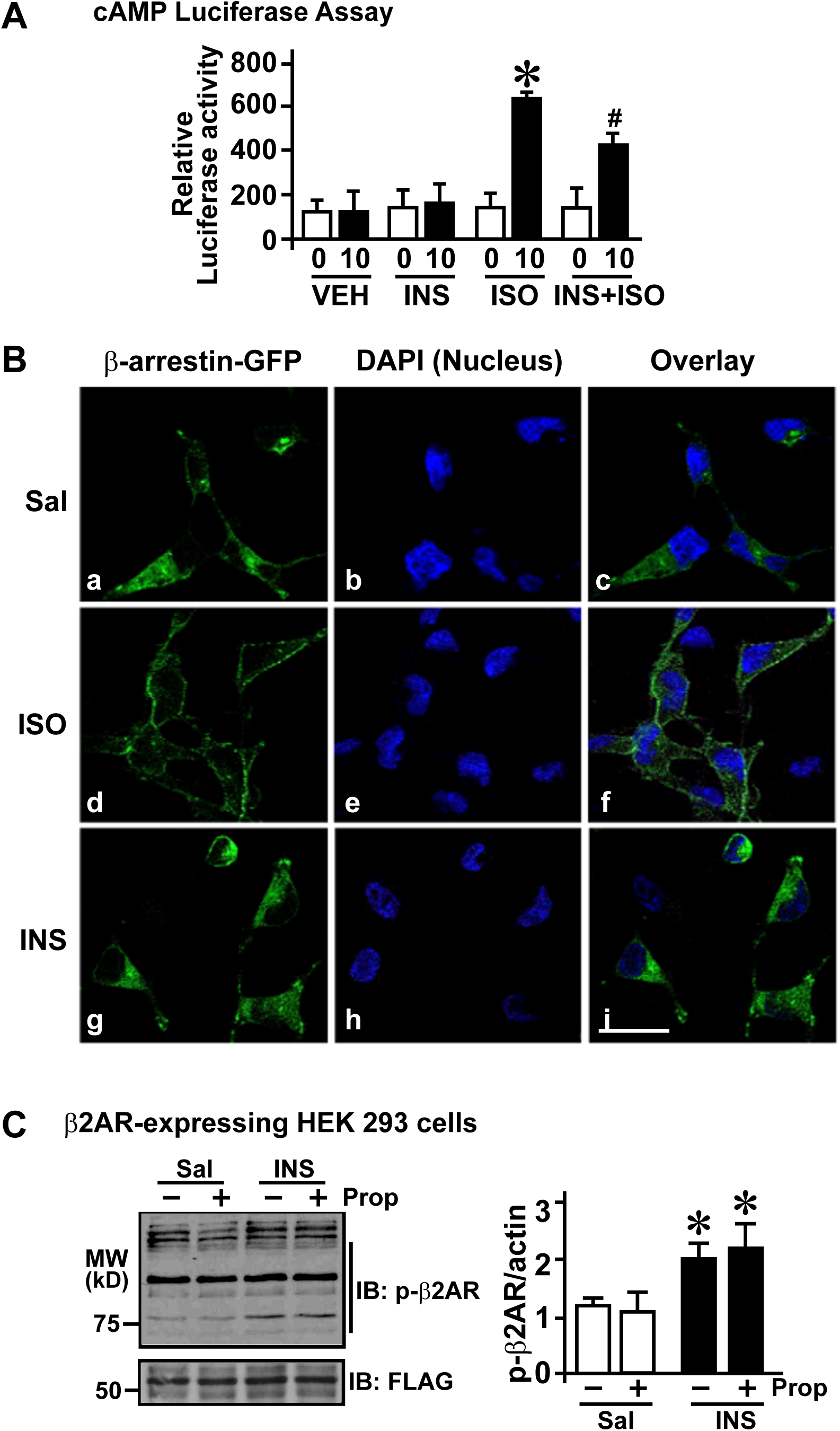
Non-canonical regulation of β2AR by INS mediates β2AR dysfunction that is β-arrestin and β-blocker independent: (A) cAMP generation was assessed using luciferase assay following pre-treatment with INS and re-challenged with ISO in FLAG-β2AR HEK 293 cells compared to untreated control cells, *p< 0.001 versus control (Sal), #p< 0.001 versus ISO. (B) β-arrestin (green) recruitment to the plasma membrane was visualized by confocal microscopy using a double stable cell line expressing β2AR and β-Arrestin 2-GFP following Sal, ISO or INS treatment for 10 minutes. Nucleus was visualized by DAPI (blue) staining. Scale bar: 10 μm (C) Immunoblotting for phospho-β2AR upon pre-treatment with propranolol (β-blocker) followed by INS treatment in the FLAG-β2AR HEK 293 cells. FLAG immunoblotting was performed as loading control. Cumulative densitometry shown as bar graph below. (n=5), *p< 0.001 versus control (Sal).

As β-blockers deter classical agonist-mediated β2AR phosphorylation and downstream cAMP signaling^20^, we investigated whether β-blocker would block non-canonical INS-mediated β2AR phosphorylation. FLAG-β2AR HEK cells were pretreated with β-blocker (propranolol) followed by INS and β2AR phosphorylation was assessed by immunoblotting. While INS treatment leads to β2AR phosphorylation, propranolol was not able to block INS-mediated β2AR phosphorylation [**Fig. 2C, cumulative bar graph (n=5)**]. These findings suggest that INS mediates β2AR phosphorylation that cannot be blocked by traditional β-blockers indicating unique but yet to be determined non-classical mechanisms of receptor desensitization.

### Insulin induces GRK2 and PI3Kγ plasma membrane translocation

Because GRK2 plays a key role in β2AR phosphorylation following INS^21^ treatment and PI3Kγ inhibits β2AR dephosphorylation, we tested whether INS stimulation leads to recruitment of GRK2 and PI3Kγ to the plasma membrane. FLAG-β2AR HEK cells were treated with INS and plasma membranes were immunoblotted with anti-GRK2 or anti-PI3Kγ antibody. Significant recruitment of GRK2 [**Fig. 3A, upper panel; cumulative bar graph left (n=5)**] and PI3Kγ [**Fig. 3A, middle panel; cumulative bar graph right (n=5)**] were observed in the plasma membranes following INS stimulation.

**Figure 3:**
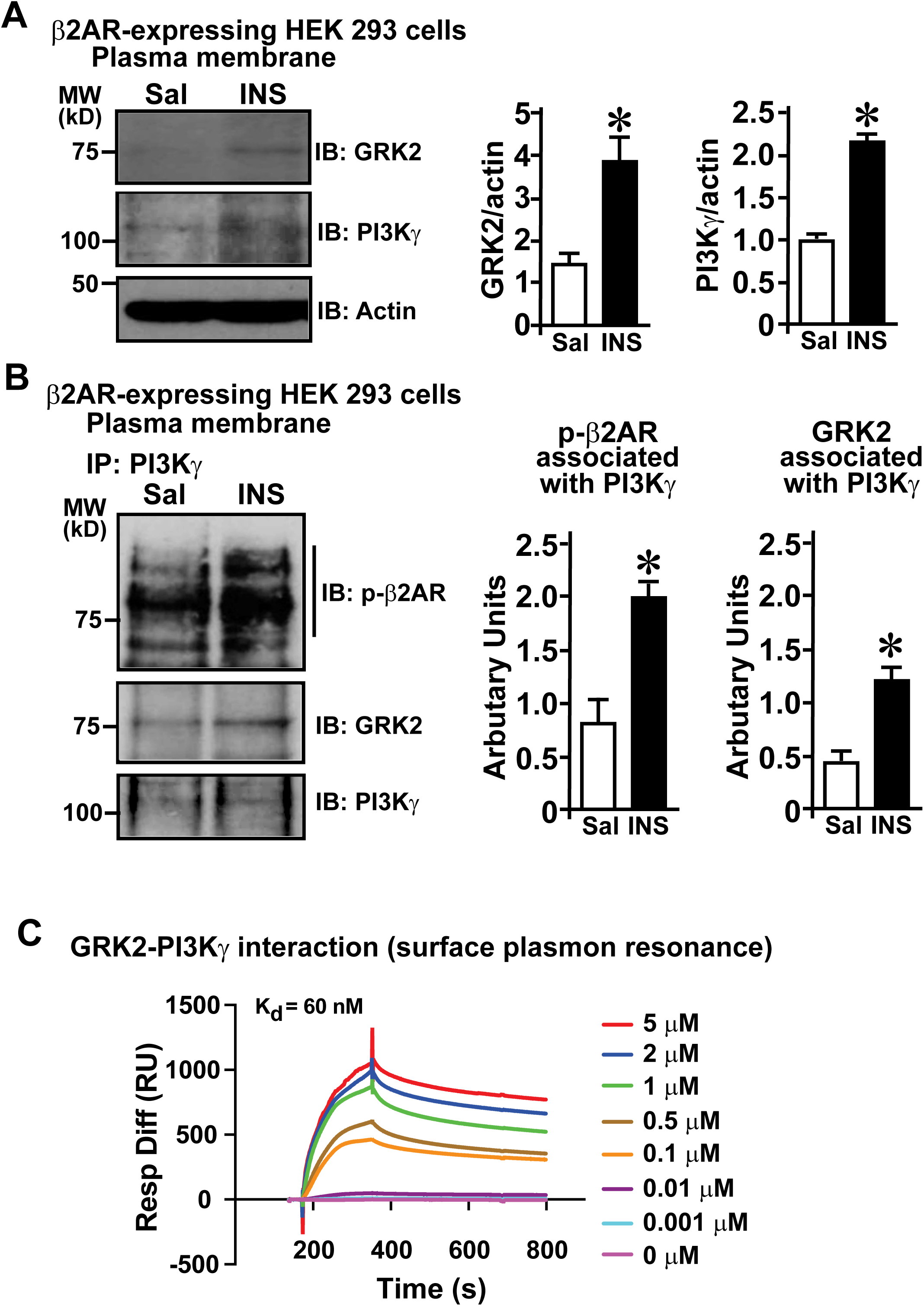
Insulin induces GRK2 and PI3Kγ plasma membrane translocation: (A) Immunoblotting for GRK2 and PI3Kγ on the plasma membranes of FLAG-β2AR HEK 293 cells following INS stimulation for 10 minutes. Cumulative data for GRK2 and PI3Kγ recruitment. (n=5), *p< 0.001 versus control (Sal). (B) PI3Kγ was immunoprecipitated from plasma membranes of FLAG-β2AR HEK 293 cells stimulated with INS for 10 minutes. Immunoprecipitates were immunoblotted for co-immunoprecipitating phospho-β2AR and GRK2. Densitometric analysis is shown in the bar graph. (n=4). *p < 0.05 compared to control (Sal). (C) Sf 9 baculovirus purified GRK2 protein was immobilized on the CM5 Biacore chip and increasing concentrations of Sf 9 baculovirus purified PI3Kγ^17^ was used as analyte. Measurement of dissociation constant shows robust interaction between GRK2-PI3Kγ with a Kd of 60 nM.

GRK2 is known to interact with IRS1/2 following INS stimulation leading to non-canonical recruitment of GRK2 to the IR-β2AR complex^13^. Consistent with these observations, immunoprecipitation of IRS1/2 showed significant interaction with GRK2 as assessed by co-immunoprecipitation [**Supplementary Fig. 1C**]. Interestingly, there was mild and appreciable co-immunoprecipitation of PI3Kγ with IRS2 immunoprecipitates but not as robust as with GRK2 [**Supplementary Fig. 1C**]. As we have previously shown that GRK2 interacts with PI3Kγ ^9^, we tested whether PI3Kγ interaction with GRK2 would be the conduit for PI3Kγ translocation to the plasma membrane following INS. PI3Kγ was immunoprecipitated from plasma membrane fraction of FLAG-β2AR HEK cells and immunoblotted for GRK2 and phosphorylated β2AR.

PI3Kγ immunoprecipitation showed significant interaction with GRK2 and phosphorylated β2AR following INS treatment [**Fig. 3B, cumulative bar graphs (n=4)**]. This shows that PI3Kγ interacts with GRK2 and is recruited to the IR-β2AR complex following INS. Consistent with this idea, immunoblotting for GRK2 in the plasma membranes from the PI3Kγ KD cells showed recruitment of GRK2 in response to INS treatment [**Supplementary Fig. 1D**]. This further supports the idea that GRK2 interaction with IRS1/2 facilitates non-canonical recruitment of PI3Kγ to the membrane following INS.

To provide unequivocal evidence that GRK2 directly interacts with PI3Kγ, we used purified proteins to assess their interaction through high sensitivity surface plasmon resonance assay. Purified GRK2 protein was immobilized on the chip and purified PI3Kγ protein was run as an analyte with increasing concentration from 0.001 μM to 5 μM. Assessment of kinetic rates of association and dissociation showed a strong protein-protein interaction between GRK2-PI3Kγ with a Kd of 60 nM [**Fig. 3C**]. These findings show that GRK2 interacts with PI3Kγ and recruits PI3Kγ to the IR-β2AR complex following INS mediating β2AR phosphorylation and dysfunction.

### PI3Kγ regulates phosphatase activity at plasma membrane after insulin stimulation

To test whether INS stimulation results in inhibition of β2AR-associated phosphatase activity, FLAG-β2AR HEK cells were stimulated with INS. FLAG-β2ARs were immunoprecipitated from plasma membranes and associated phosphatase activity was assessed by malachite green assay. FLAG-β2AR-associated phosphatase activity was significantly lower in INS treated cells compared to Sal controls [**Fig. 4A, cumulative bar graph (n=5)**]. In contrast, FLAG-β2AR phosphatase activity was normalized (*ie*, not inhibited) in the PI3Kγ KD cells [**Fig. 4B, cumulative bar graph (n=5)**] showing that non-canonical recruitment of PI3Kγ to the β2AR complex mediates inhibition of PP2A. Given that we have previously shown that PI3Kγ inhibits PP2A by phosphorylating endogenous inhibitor of PP2A, I2PP2A, we assessed whether non-canonical recruitment of PI3Kγ can mediate I2PP2A phosphorylation using a novel in-house custom-made phospho-I2PP2A antibody (see methods for details). Significant phosphorylation of I2PP2A was observed in Sh vector control cells and was completely abrogated in the PI3Kγ KD cells [**Fig. 4C, cumulative bar graph (n=5)**].

**Figure 4:**
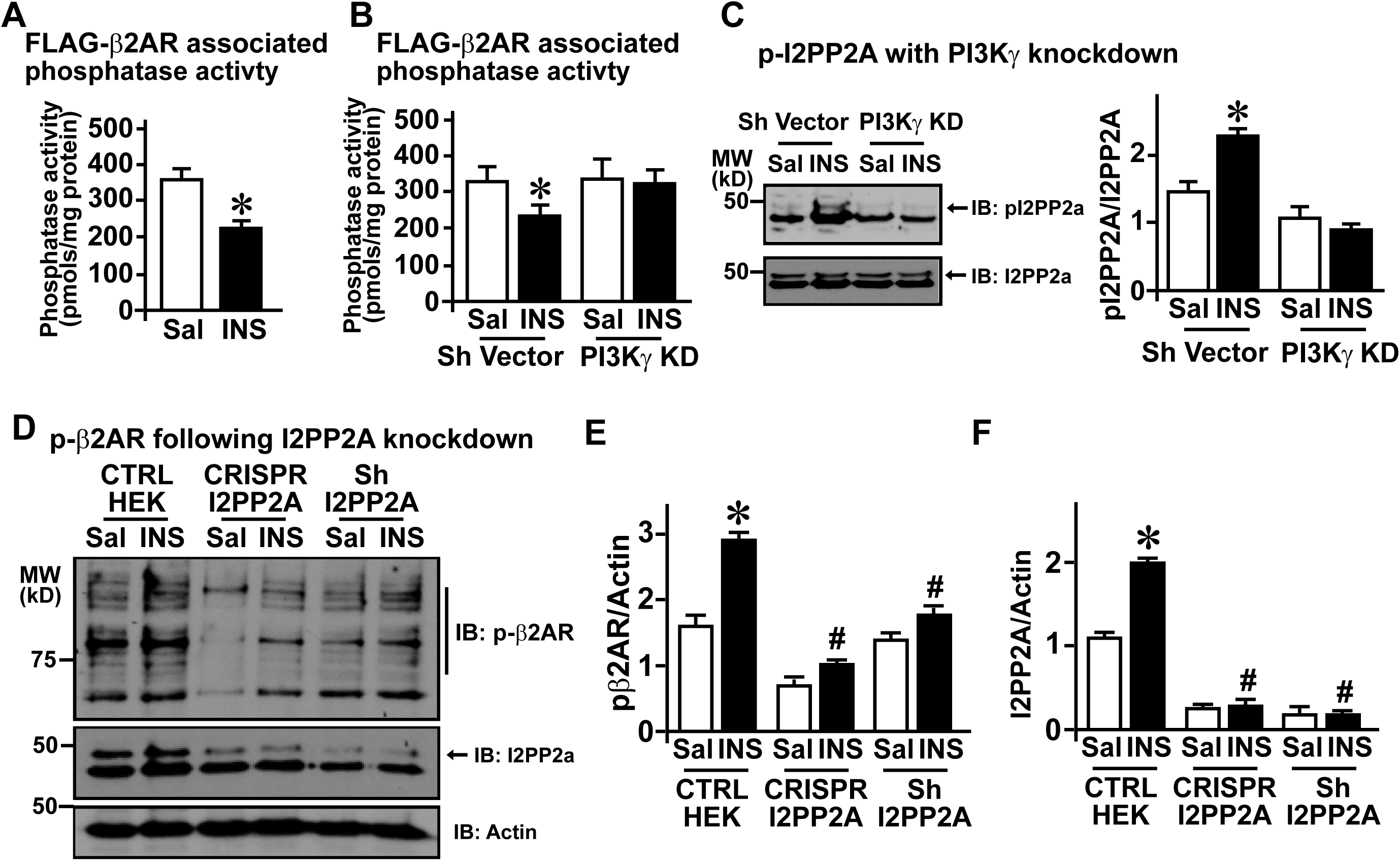
PI3Kγ regulates phosphatase activity at plasma membrane after Insulin stimulation. (A) FLAG-β2AR immunoprecipitated from plasma membrane fraction of FLAG-β2AR HEK 293 cells and associated phosphatase activity assessed following INS stimulation for 10 minutes. (n=3). *p < 0.001 compared to control (Sal). (B) FLAG-β2AR was transfected into Sh Vector and PI3Kγ KD HEK 293 cells and FLAG-β2AR immunoprecipitated from plasma membranes and assessed for associated phosphatase activity following INS stimulation for 10 minutes, (n=3). *p < 0.001 compared to control (Sal). (C) Immunoblotting was performed to assess phospho-I2PP2A levels in Sh Vector and PI3Kγ KD cells following INS stimulation for 10 minutes. Total I2PP2A was immunoblotted as loading control. Cumulative densitometry is presented in the bar graphs. (n = 5). *p < 0.001 compared to control (Sal). (D) FLAG-β2AR was transfected into HEK 293 parental cells, CRISPR I2PP2A or stable Sh I2PP2A knock down cells and immunoblotting was performed to assess phospho-β2AR, phospho-I2PP2A and I2PP2A in total cell lysates. Cumulative densitometry is presented in the bar graphs. (n = 4). *p < 0.001 compared to control (Sal).

To determine whether I2PP2A inhibits PP2A and plays a key role in maintaining phosphorylation of β2AR following INS, we generated HEK 293 cells with CRISPR-mediated I2PP2A ablation and stable Sh-RNA-mediated knockdown of I2PP2A. These cells were transfected with FLAG-β2AR expressing cDNA construct and stimulated with INS followed by assessment of β2AR phosphorylation by immunoblotting. INS treatment consistently resulted in increased β2AR phosphorylation in control HEK 293 cells which was significantly abrogated in both the I2PP2A knockdown cells [**Fig. 4D, cumulative bar graphs (n=4)**]. These findings show the key role of PI3Kγ mediated inhibition of PP2A through phosphorylation of I2PP2A that sustains β2AR phosphorylation following INS stimulation.

### PI3Kγ regulation of INS-mediated β2AR phosphorylation is conserved in primary cardiac cells

As a first step to test whether INS mediates β2AR phosphorylation in vivo, mice were administered Sal or INS for two weeks. Hearts were excised and total cardiac lysates were immunoblotted with anti-phospho-β2AR antibody to assess receptor phosphorylation. INS administration resulted in significant increase in β2AR phosphorylation compared to Sal treatment [**Fig. 5A, cumulative bar graph (n=6)**]. To further test whether phosphorylation of β2ARs occur in cardiomyocytes, primary cardiomyocytes were isolated from mice hearts following two weeks of INS administration and assessed for β2AR phosphorylation, PI3Kγ and GRK2 levels. Significant increase in a β2AR phosphorylation was observed in cardiac myocytes from INS administered mice [**Fig. 5B**] and was associated with mild increase in GRK2 and PI3Kγ levels [**Fig. 5B, lower panels**] which did not attain the level of significance. Given that our cellular studies have focused on acute INS regulation of β2AR phosphorylation by PI3Kγ, we performed acute in vivo and ex vivo cellular studies. Mice were acutely administered with INS acutely for 20 minutes and total cardiac lysates from the excised hearts were assessed for β2AR phosphorylation. Interestingly, acute challenge of INS results in significant increase in β2AR phosphorylation compared to Sal [**Fig. 1C, cumulative bar graph (n=6)**]. To further dissect whether PI3Kγ regulates β2AR phosphorylation in response to INS, primary adult cardiomyocytes and primary cardiac fibroblasts were isolated from wild type (WT) and PI3Kγ knockout (KO) mice. Primary cardiomyocytes were treated with INS for varying period of time (0 – 20 minutes) after 4 hours of plating the isolated cells and the lysates were immunoblotted with anti-phospho-β2AR antibody. INS treatment resulted in significant phosphorylation of β2AR in WT adult cardiomyocytes which was remarkably abolished in PI3Kγ KO cardiomyocytes [**Fig. 1D, cumulative bar graph (n=5)**]. Primary cardiac fibroblasts were grown for a week, serum starved and acutely stimulated with INS for 10 minutes.

**Figure 5:**
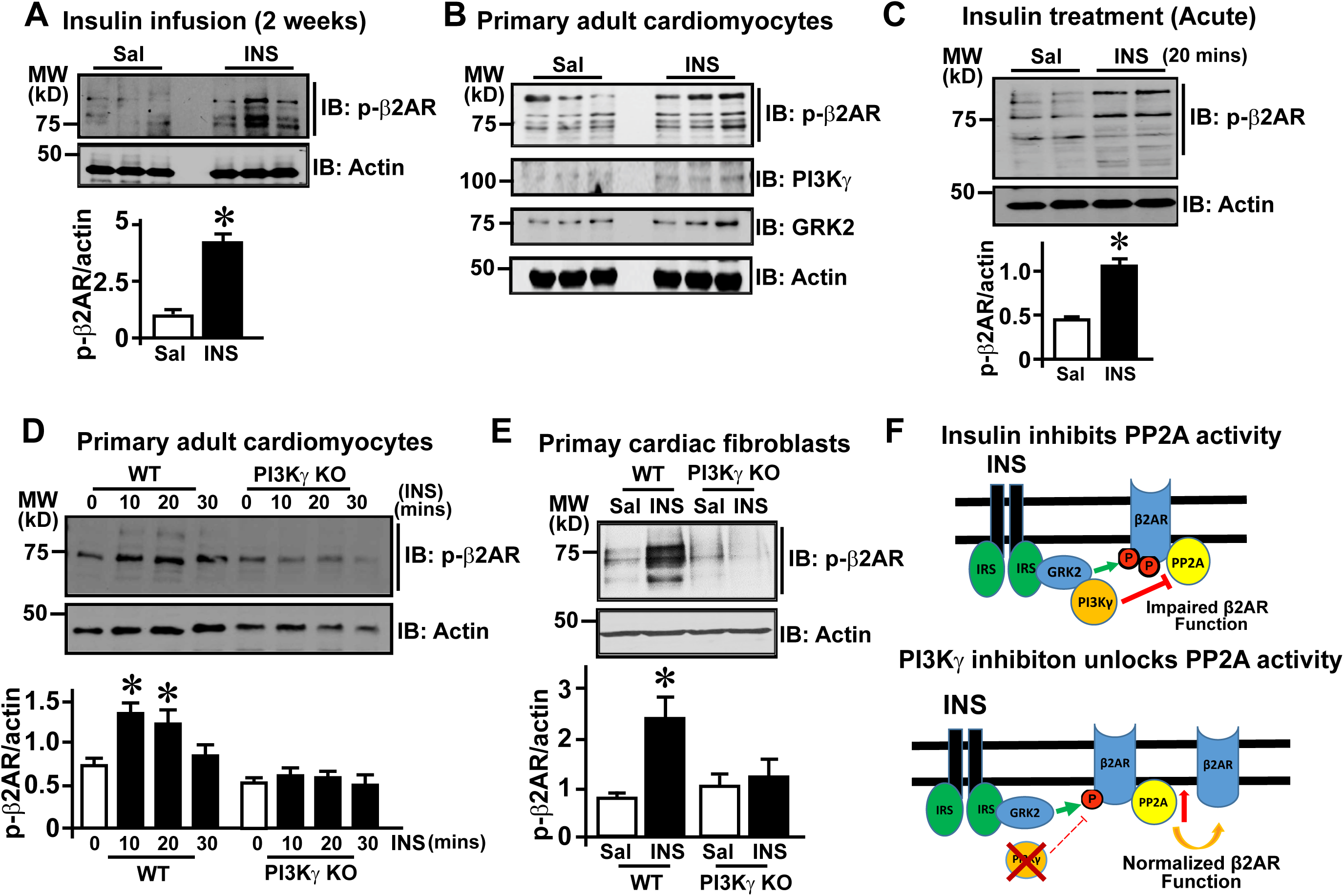
PI3Kγ regulation of INS-mediated β2AR phosphorylation is conserved in primary cardiac cells: (A) Immunoblots to assess phospho-β2AR in cardiac lysates from mice infused with Sal or INS for 2 weeks. Cumulative data is presented in the bar graphs. (n = 6). *p < 0.0001 compared to control (Sal). (B) Immunoblotting for phospho-β2AR, GRK2 and PI3Kγ in total cell lysates of primary cardiac myocytes isolated from mice infused with Sal or INS for 2 weeks. (n=6). (C) Immunoblots to assess phospho-β2AR in cardiac lysates from mice injected with Insulin or Saline for 20 minutes. Densitometry is presented in the bar graphs. (n = 6). *p < 0.0001 compared to control (Sal). (D) Immunoblots to assess phospho-β2AR in primary cardiac myocytes isolated from WT and PI3Kγ KO mice and stimulated with INS for 0, 10, 20 and 30 minutes. Cumulative data is presented in the bar graphs. (n = 5). *p < 0.001 compared to control (Sal). (E) Immunoblots to assess phospho-β2AR in primary cardiac fibroblasts isolated from WT and PI3Kγ KO mice and stimulated with INS for 10 minutes. Cumulative densitometry is presented in the bar graphs. (n = 5). *p < 0.001 compared to control (Sal).

Immunoblotting with anti-phospho-β2AR antibody showed significant phosphorylation in the WT cardiac fibroblasts to INS, which was completely abrogated in the PI3Kγ KO cardiac fibroblasts [**Fig. 5E, cumulative bar graph (n=5)**]. Together these studies show that regulation of INS mediated phosphorylation of β2AR by PI3Kγ is conserved in the heart as well as in the adult cardiomyocytes and fibroblasts.

## Materials and methods

### Animals

C57/BL6 mice null for PI3Kγ (PI3Kγ knock out, PI3Kγ KO (described previously^1^) and C57/BL6 wild type (WT) mice were administered insulin (INS) (0.1 IU/g/day for 14 days (chronic) or 1 IU/kg for 20 min (acute)). The mice were euthanized, and hearts excised for biochemical analysis. All experiments involving animals were performed on approved protocol and in accordance with institutional guidelines and regulations of IACUC/Institutional Review Board at Cleveland Clinic.

### Pharmacological compounds

Isoproterenol hydrochloride, Insulin, Propranolol, LY294002, and wortmannin were all obtained from Sigma-Aldrich. In addition, all the chemicals used for making buffers have been purchased from Sigma-Aldrich or ThermoFisher and if not are mentioned as they are listed in the methods section.

### Cell culture (HEK 293 and mouse embryonic fibroblasts)

#### HEK 293 cells

HEK 293 parental cells or HEK 293 cells stably overexpressing FLAG-β2AR (FLAG-β2AR HEK), stable knockdown of I2PP2A (I2PP2A KD)^7^ or PI3Kγ (PI3Kγ KD)^17^ were used in the study. Sequences and details of the siRNAs used, and development of stable shRNA knockdown cells have been described previously^9^. These cell lines were grown and maintained in MEM (Life Technologies) with 10% FBS (Life Technologies) and 5 % penicillin/streptomycin (Life Technologies). I2PP2A KD or PI3Kγ KD HEK293 cells were also maintained in MEM with 10% FBS supplemented with hygromycin (400 mg/ml).

#### Mouse embryonic fibroblast (MEF)

Isolation and maintenance of MEFs from wild-type and PI3Kγ knockout mice have been described previously^22^. MEFs were maintained in MEM with 10% FBS and 5%penicillin/streptomycin. All cells were maintained in a humidified incubator at 37°C under 5% CO2. “n” represents the number of independent experiments performed.

#### Treatment and Transfection

HEK 293 cells or FLAG-β2AR HEK 293 cells in serum free media (4 hours) were treated with insulin (INS) (100nM), isoproterenol (ISO) (10 μM) or Propranolol (PROP) (10 μM) either for 10 minutes for different time points to assess phosphorylation of β2AR. Furthermore, cells were pretreated with Wortmannin (Wort) (100 nM) or LY29002 (LY) (1 μM) separately or along with INS. Also, HEK 293 cells with stable knockdown of I2PP2A (I2PP2A KD) or PI3Kγ KD were transfected with 2 μg of FLAG-β2AR expressing plasmid construct using Turbofect (ThermoFisher) following the manufacturer’s protocol. 48-hours post-transfection, the cells were serum starved and treated with INS. Controls in the experiments underwent similar conditions except that they were treated with saline (Sal).

#### CRISPR construct and sgRNA transfection

CRISPR sgRNAs were generated from human I2PP2A cDNA sequence using design programs (http://crispor.org). The target 23-mer sequences for sgRNA used were 5’-TTTACCTTGATAGGTGGTAGTGG - 3’and 5’-ATAAAGAGAGGCTTCTCGGGAGG - 3’ and synthesized using Guide-it™ sgRNA In Vitro Transcription Kit (TaKaRa) according to manufacturer’s protocol. One day prior to transfection, HEK 293 cells were plated onto 6-well plates (2.5-4.5 × 10^5^ cells per well) in 3 ml of MEM (Life Technologies) with 10% FBS (Life Technologies) and 5%penicillin/streptomycin (Life Technologies). Cells were transfected with optimum concentration of sgRNAs using Lipofectamine™ CRISPRMAX™ Cas9 Transfection Reagent (ThermoFisher) following the manufacturer’s protocol. 48–72 hours post-transfection, the cells were harvested for analysis using Guide-it™ sgRNA Screening Kit (TaKaRa). Stable I2PP2A knockdown CRISPR I2PP2A HEK293 cells were also maintained in MEM with 10% FBS supplemented with Gentamycin (50 mg/ml).

#### Plasma membrane isolation

Plasma membranes were isolated as described previously^23^. Briefly, cells were lysed in ice-cold lysis buffer containing 5 mM Tris-HCl pH 7.4, 5 mM EDTA, 1 mM PMSF, and 2 μg/mL leupeptin and aprotinin. Intact cell debris and nuclei were removed by centrifugation at 2,500 X *g* for 5 minutes and the supernatant was subjected to centrifugation at 37,000 X *g* for 20 minutes. The pelleted plasma membrane was resuspended in triton buffer (20 mM Tris-HCl, pH 7.4, 300 mM NaCl, 1.0 % triton X-100, 1 mM PMSF, 20% glycerol, 10 mM NaF, 1 mM sodium orthovanadate, 2 μg/mL leupeptin and aprotinin) for the immunoprecipitation experiment or NP-40 lysis buffer [20 mM Tris, pH 7.4, 137 mM NaCl, 1 mM phenylmethanesulfonyl fluoride (PMSF), 20% glycerol, 10 mM NaF, 1% NP-40, sodium orthovanadate, leupeptin, aprotinin, phosphatase inhibitor cocktail] for western immunoblotting studies.

#### Western immunoblotting

Western immunoblotting was performed as described previously^9^. Briefly, cell lysates (50 - 100 μg) or plasma membrane fractions (80 - 100 μg) were resolved on SDS-PAGE gel and transferred to PVDF membrane (BIORAD). The membranes were blocked with 5% BSA and immunoblotted using primary anti-phospho-β2AR antibody (1:1000) (Santa Cruz Biotechnology), anti-PI3Kγ (1:500) (Santa Cruz Biotechnology), anti-GRK2 (1:1000) (Santa Cruz Biotechnology), anti-I2PP2A (1:700) (Santa Cruz Biotechnology or) or anti-PP2Ac (1:3000) (Santa Cruz Biotechnology).

#### Generation of phospho-I2PP2A antibody

Anti-phospho-I2PP2A anti-body was developed in-house at the Cleveland Clinic using the hybridoma core facility. Polyclonal antibody against the phosphorylated form of I2PP2A was generated by immunizing rabbits with a synthetic peptide corresponding to residues surrounding Ser9 of human I2-PP2A (APAAKCpSKKELNC) as previously described^9^. The specificity of the antibody was tested by immunoblotting in vitro phosphorylated purified I2-PP2A (phospho-I2PP2A) protein and in lysates from isoproterenol-stimulated HEK 293 cells. The purified protein used in the assay to authenticate the antibody was validated using mass spectrometry in proteomics core facility. Following immunoblotting with primary antibody, horseradish peroxidase-conjugated secondary antibodies (GE Healthcare or Thermo Scientific) were used for detection using chemiluminescence. Image analysis was performed with National Institutes of Health (NIH) ImageJ software.

#### Phosphatase assay

Phosphatase activity was assessed using the serine-threonine phosphatase assay kit (Cat#20-105; Upstate Biotechnology). Samples of 200 μg of membrane protein were used for FLAG immunoprecipitation. Following immunoprecipitation, the samples were resuspended in the phosphate-free assay buffer and incubated in the presence or absence of serine-threonine specific phosphopeptide substrate for 10 min. Purified PP2A supplied by the manufacturer was used as a positive control to normalize the activity in the samples. The reaction mix was incubated with acidic malachite green solution, and absorbance at 630 nm was measured in a plate reader.

#### cAMP Luciferase assay

FLAG-β2AR HEK 293 cells were transfected with 3 µg cAMP luciferase plasmid DNA (Promega) using Turbofect (ThermoFisher) following the manufacturer’s protocol. 24 hours post-transfection, the cells were trypsinized and plated onto a 96-well plate at 50,000 cells/well and allowed to settle for 12 hours. The cells were stimulated with INS (100nM) or ISO (10uM) and the cAMP generation was measured according to manufacturer’s instruction using the GloSensor™ (Promega) assay system by measuring the luciferase activity **us**ing Molecular Probes measurement platform.

#### Confocal microscopy

FLAG-β2AR HEK 293 cells or HEK 293 cells stably expressing β2AR and β-arrestin 2-GFP (generous gift from Dr. Robert Lefkowitz) were plated on to cover slips treated with poly L-lysine and serum starved for 4 hours prior to ISO or INS treatment. Since ISO treatment results in marked endocytosis of β2ARs, we used endocytosis inhibitor (0.45M sucrose and 2% Cyclodextrin for 1 hour) so that we could visualize β2AR phosphorylation on the plasma membrane. Since we do not know whether INS causes receptor internalization acutely, cells that were to be stimulated with INS were not treated with endocytosis inhibitors. After ISO or INS stimulation, cells were fixed (4% paraformaldehyde), permeabilized with ice cold methanol for 10 min at –20°C and blocked with 5% normal goat serum (NGS) in PBS for 1 hour. Anti-phospho-β2AR antibody (1:1000; Santa Cruz) incubation (overnight) was followed by secondary goat anti-rabbit AlexaFlour 488 conjugated antibody (1:200; Molecular Probes). HEK 293 cells stably expressing β2AR and β-arrestin 2-GFP were stimulated with ISO or INS for 10 minutes and were fixed for visualization. Samples were visualized using sequential line excitation at 488 for green and nuclei was visualized by DAPI using Leica TCS-SP8-AOBS inverted confocal microscope using LAS X v3.1.5 with HyD detectors (Leica Microsystems, GmbH).

#### Isolation of primary adult cardiomyocytes and primary cardiacfibroblasts

The hearts were excised out of anesthetized mice, and immediately cannulated with 20-gauge needle and perfused with collagenase. Hearts were then perfused with cold phosphate-buffered saline (PBS) before being minced into 1–2 mm pieces and centrifuged for 3 min at 20 x g to separate out small non-myocyte cells such as endothelial cells and fibroblasts (supernatant) from cardiomyocytes (pellet). The myocyte pellet was resuspended in a plating medium of MEM containing 10% FBS, 500mM 2, 3-butanedione monoxime (BDM) and 200 mM ATP. The cells were plated on a 60-mm culture dish and incubated for 2 h at 37°C in a humidified incubator under 2% CO2. The primary cultures of cardiac myocytes were maintained in the medium MEM, 100 mg/ml BSA, and 2.2% antibiotic/antimycotic (10,000 U/ml penicillin, 10,000 μg/ml streptomycin, and 25 μg/ml amphotericin B; Life Technologies). Supernatants collected in a fresh tube were centrifuged at 1000 rpm for 5 minutes and the pellet was resuspended in MEM containing 10% FBS, plated on a 100-mm culture dish, and incubated for 2 h at 37°C in a humidified incubator under 5% CO2. Following 2 hours of incubation, the unattached cells were discarded and attached cells were grown in MEM containing 10% FBS and 2.2% antibiotic/antimycotic (10,000 U/ml penicillin, 10,000 μg/ml streptomycin, and 25 μg/ml amphotericin B; Life Technologies) at 37°C. The attached cells are primary cardiac fibroblasts and were maintained in a humidified incubator at 37°C and under 5% CO2 and the medium was changed every 2 days. Primary cardiac fibroblasts were stimulated with INS after 7 days of culture.

#### Surface Plasmon Resonance assay

Biomolecular interaction analysis was performed using a Biacore 3000 (GE Healthcare Life Sciences) in the Molecular Biotechnology Core of Lerner Research Institute, Cleveland Clinic. Binding was measured by recording response units (RU) over time. Purified GRK2^24^ was immobilized on CM5 Sensor Chip (GE Healthcare Life Sciences) using 10 mM acetate buffer, pH 5.5. Purified 6-His-PI3Kγ catalytic subunit^17,18^ was used as the soluble analyte and was injected at different concentrations over the immobilized GRK2 in SPR buffer (0.01 M HEPES pH 7.4, 0.15 M NaCl, 3 mM EDTA, 0.005% surfactant P-20, GE Healthcare Life Sciences, Biacore Inc.). Binding experiments were performed at 25°C in SPR buffer with a flow rate of 20 μl/min. Data was normalized against an empty reference channel. Analysis and fitting were performed with BIAevaluation software, version 4.0.1 (Biacore Inc.), with the option for calculating the association/dissocation constant (Ka/Kd). Global fitting was used for fitting sensogram data, and a 1:1 Langmuir model was used to calculate the Ka and Kd values. The distribution and magnitude of the residuals, as well as χ2 values, were used to determine fit quality^25^.

#### Statistical analysis

All data expressed as mean ± SEM (*n* ≥ 3). Analysis of variance was used for multiple comparisons of the data. Statistical analyses were performed using GraphPad Prism, and the significance between the treatments was determined by Student’s *t* test. A *p* value less than 0.05 was considered statistically significant.

## Discussion

Cross talk between β2AR and receptor tyrosine kinases like EGFR, IGFR/IR and PDGFR is well known^26^. β2AR activation results in transactivation of receptor tyrosine kinase EGFR leading to activation of downstream signals^27^. However, activation of IGFR/IR by insulin mediates desensitization of β2AR, impairing β2AR function^10^ that is known to be regulated by GRK2-dependent mechanisms whereas, less is known about dephosphorylation mechanisms. In our current study, we show that acute regulation of phosphatase by INS stimulation plays as important a role in sustaining kinase-driven impairment of β2AR function. Interestingly, inhibition of PI3K or specifically knock down or ablation of PI3Kγ results in complete reversal/abrogation of β2AR phosphorylation in response to INS. This shows a unique regulation of receptor cross-talk by PI3Kγ that is supported by studies in primary adult cardiomyocytes and cardiac fibroblasts reflecting the existence of a conserved under-appreciated regulation of IR and β2AR cross-talk by PI3Kγ. Using co-immunoprecipitation studies and purified proteins we show that PI3Kγ interacts with GRK2 allowing for non-canonical recruitment of PI3Kγ to the functional IR-β2AR complex through GRK2-IRS interaction axis. INS stimulation inhibits β2AR-associated PP2A activity in part, accounting for sustained phosphorylation of β2AR as knock down of PI3Kγ unlocks the inhibition of β2AR-associated PP2A activity. This unlocking results in loss of β2AR phosphorylation in spite of INS. Also, INS stimulation led to phosphorylation of the endogenous inhibitor of PP2A (I2PP2A) whose knock down reverses INS-mediated β2AR phosphorylation showing that acute inhibition of PP2A is an integral mechanism in sustaining INS-mediated receptor phosphorylation. These findings show a pivotal role for PP2A regulation by PI3Kγ which is recruited non-canonically to the IR-β2AR complex via its interaction with GRK2.

Traditionally, PI3Kγ is recruited to the β2AR complex following agonist-stimulation by the dissociated Gβγ subunits of G-proteins^28^. Recruitment of PI3Kγ by Gβγ subunits initiates PI3K dependent downstream signaling following agonist activation of β2AR^29^. Further, recruitment of PI3Kγ also plays a key role in β2AR-mediated EGFR transactivation by regulating non receptor tyrosine kinase Src^16^. However, in contrast to this established mechanism of PI3Kγ recruitment to the plasma membrane, INS stimulation *per se* does not result in G-protein activation or dissociation of G-protein subunits suggesting non-canonical recruitment of PI3Kγ to the receptor complex. Consistent with our previous study^30^, we have found that PI3Kγ interacts with GRK2 to form a complex. In addition, immunoprecipitation of PI3Kγ from the plasma membranes showed significant interaction between GRK2 and phosphorylated β2AR following INS showing a key role for GRK2-PI3Kγ in regulating β2AR function. Importantly, using purified proteins we unequivocally show that the interaction between GRK2 and PI3Kγ is very robust (Kd of 60 nM) allowing for Gβγ-independent recruitment of PI3Kγ to the functional IR-β2AR complex through GRK2. This idea is supported by co-immunoprecipitation findings [**Supplementary Fig 1C**] and previous studies showing robust interaction between GRK2 and IRS 1/2^14^. IRS2 interaction with GRK2 is required for GRK2 recruitment to the IR-β2AR complex as absence of IRS2 results in loss of INS-mediated β2AR phosphorylation^13^.

These findings suggest that the GRK2-PI3Kγ recruitment to the IR-β2AR functional complex mediates a unique and yet sustained dysregulation of β2AR function. It is unique given that GRK2 mediates β2AR phosphorylation even in the absence of its classical agonist and this non-canonical receptor phosphorylation is maintained/sustained by inhibition of β2AR-associated PP2A activity by PI3Kγ. Acute inhibition of PP2A activity by non-canonical recruitment of PI3Kγ sustains GRK2 mediated phosphorylation of β2AR as inhibition of PI3K or specifically knock down of PI3Kγ unlocks this inhibition leading to abrogation of INS-mediated β2AR phosphorylation. This is supported by our observation that PI3Kγ KD completely reverses β2AR-associated phosphatase activity even in the presence of INS accounting for effective dephosphorylation of β2AR. Importantly, there was no inhibition of ISO-stimulated cAMP generation following INS pre-treatment in PI3Kγ KD cells reflecting that dephosphorylation of β2ARs now allows for active engagement of the agonist. Moreover, significant loss in β2AR phosphorylation is also observed in primary adult cardiomyocytes and cardiac fibroblasts isolated from PI3Kγ KO mice. This shows that the unique PI3Kγ-mediated regulation of PP2A occurs in vivo and mechanistic understanding of this underappreciated pathway may provide therapeutic strategies for preserving β2AR function in hyperinsulinemia.

PI3Kγ, as a member of the PI3K family is an established lipid kinase when recruited to the plasma membrane utilizes phosphoinositide-4,5-*bis*-phosphate (Ptd-4,5-PO4) to go generate Ptd-3,4,5-PO4 that now initiates downstream Akt signaling^31^. However, we have previously shown that protein kinase activity of PI3Kγ regulates PP2A function at the β2AR complex following agonist stimulation^9^. In this context, acute regulation of PP2A function is mediated by phosphorylation of an endogenous inhibitor of PP2A (I2PP2A) by PI3Kγ that binds robustly to PP2A inhibits PP2A activity sustaining βAR phosphorylation following agonist stimulation. Consistent with these studies, non-canonical recruitment of PI3Kγ mediates I2PP2A phosphorylation following INS (as assessed by our newly generated in-house phospho-I2PP2A antibody) which was completely abolished with PI3Kγ KD despite the presence of INS. These observations provide evidence that acute PP2A inhibition by PI3Kγ is as critical a player in mediating and sustaining β2AR phosphorylation as GRK2. Such a concept is supported by the evidence that GRK2-mediated phosphorylation becomes inconsequential when β2AR-associated PP2A activity is preserved without inhibition. This is further supported by the observation with siRNA or CRISPR knock down of I2PP2A which disrupts PI3Kγ-mediated inhibition of PP2A. Thus, releasing this inhibitory regulation results in marked loss of β2AR phosphorylation despite presence of INS, GRK2 and PI3Kγ reflecting the key role of PP2A inhibition in mediating β2AR dysfunction in the receptor cross-regulation.

It is well known that β-blockers block agonist-mediated βAR activation and G-protein coupling^32^ but by themselves can initiate G-protein independent β-arrestin-dependent beneficial signals^33^. In contrast to their role in blocking agonist-mediated β2AR activation, β-blocker pre-treatment did not inhibit INS-mediated β2AR phosphorylation. This shows that non-canonical recruitment of GRK2 to the functional IR-β2AR complex in response to INS is still able to recognize β2AR as a substrate and mediate phosphorylation even in the presence of β-blocker. This is an interesting observation given that β-blockers are prescribed for heart failure in diabetes and hyperinsulinemia^34^ indicating that β-blocker may not be as effective. Interestingly, there was also no acute recruitment of β-arrestin to the β2ARs despite phosphorylation suggesting that INS-mediated receptor regulation may not alter receptor confirmation or engage the classical agonist dependent pathways to mediate β2AR dysfunction. However, more mechanistic studies are needed to understand the underpinnings of GRK2 mediated β2AR phosphorylation despite the presence of β-blocker.

These observations show that INS non-canonically recruits GRK2-PI3Kγ to the IR-β2AR functional complex mediating acute inhibition of PP2A through I2PP2A resulting in β2AR dysfunction. Identification of this PP2A inhibitory pathway shows an additional layer in the regulation of INS-mediated β2AR dysfunction. However, this is a pivotal regulator in the INS-IR-β2AR cross-talk as disengaging the phosphatase inhibitory pathway completely preserves βAR function despite the presence of INS and GRK2. Given that increasing number of studies have shown a direct correlation between insulin resistance and heart failure^5, 35^ our studies provide novel insights into mechanisms that impair β2AR function by INS. Importantly, these studies bring-to-fore the concept that acute regulation of PP2A especially their inhibition could play a critical role in mediating and sustaining kinase driven signaling pathways.

## Acknowledgements

The studies are supported in part by AHA TPA (18TPA34170554) to MLM, RO1 HL071818 (JJGT) and RO1 HL128382 and R01 HL089473 (SVNP). Phospho-I2PP2A antibody were generated by the hybridoma core at the Lerner Research Institute, Cleveland Clinic. We would like to thank Dr. Smarajit Bandyopadhay (Biotechnology Core, Lerner Research Institute) for help in conducting SPR experiments.

## Contributions

Anita Sahu, Yu Sun, Sromona Mukherjee, Conner Witherow, Kate Stenson, Maradumane L. Mohan and Sathyamangla V. Naga Prasad: Designed, performed and optimized experiments, analyzed and interpreted data, drafted and edited the final manuscript. John J.G. Tesmer: Analyzed and interpreted the data, drafted, and edited the manuscript.

---

**Supplementary Figure 1:**
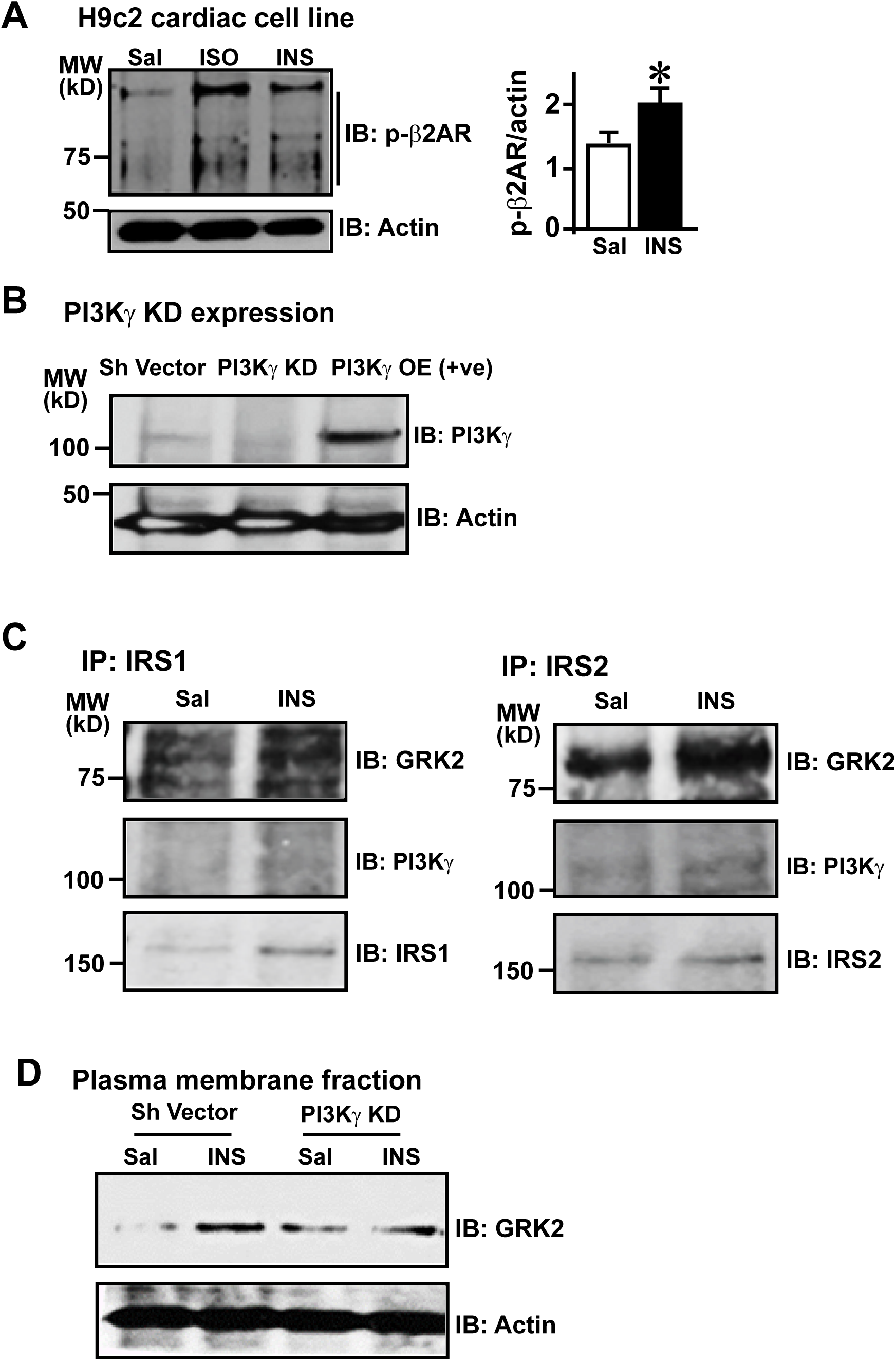
**Figure 1A** - β2AR phosphorylation in H9c2 cardiac myoblasts following INS stimulation: H9c2 cardiac myoblasts were serum starved for 4 hours followed by stimulation with classical β2AR agonist isoproterenol (ISO, 10 μM) or insulin (INS, 100 nM). H9c2 cell lysates were immunoblotted with anti-phospho-β2AR antibody to assess β2AR phosphorylation. Significant β2AR phosphorylation of endogenous β2ARs is observed following ISO or INS. Cumulative data for INS-mediated phosphorylation is presented in the bar graphs. (n = 3). *p < 0.001 vs. saline control (Sal). β-actin was used as loading control. **Figure 1B** - PI3Kγ expression level in PI3Kγ stable knockdown and overexpressed HEK 293 cells: Expression of PI3Kγ was assessed in control (Sh Vector), stable expression of Sh-RNA targeting PI3Kγ (PI3Kγ KD) and as positive control (+ve) cells stably overexpressing PI3Kγ (PI3Kγ OE). Cell lysates were immunoblotted with anti-PI3Kγ antibody. β-actin was used as loading control. **Figure 1C** - GRK2 and PI3Kγ interaction with IRS1 and IRS 2: GRK2 has previously been shown to interact with IRS1 and IRS2. To test whether GRK2, PI3Kγ and IRS forms a complex, endogenous IRS1 and IRS2 were immunoprecipitated from plasma membranes of FLAG-β2AR HEK 293 cells stimulated with INS (100 nM) for 10 minutes. Immunoblotting showed co-immunoprecipitation of GRK2 and PI3Kγ, (n=3). **Figure 1D** - Recruitment of GRK2 in response to INS stimulation in the absence of PI3Kγ: Since absence of PI3Kγ in PI3Kγ KD cells results in abrogation of β2AR phosphorylation with INS, we tested whether GRK recruitment to the plasma membrane still occurs following INS. Immunoblotting for GRK2 in plasma membranes of Sh Vector and PI3Kγ KD HEK 293 cells transfected with FLAG-β2AR following INS stimulation (100 nM) for 10 minutes. (n=4) shows robust recruitment of GRK2 to the plasma membrane in PI3Kγ KD cells and yet we see reduced β2AR phosphorylation consistent with increased β2AR-associated phosphatase activity (Fig. 4B).

## Notes

### Competing Interest Statement

The authors have declared no competing interest.

